# Unifying DNA methylation-based in silico cell-type deconvolution with *deconvMe*

**DOI:** 10.1101/2025.03.10.642382

**Authors:** Alexander Dietrich, Lina-Liv Willruth, Korbinian Pürckhauer, Carlos Oltmanns, Moana Witte, Sebastian Klein, Anke RM Kraft, Markus Cornberg, Markus List

**Affiliations:** Data Science in Systems Biology, TUM School of Life Sciences, Technical University Munich, Munich, Germany; Core Facility Microbiome, ZIEL Institute for Food & Health, Technical University of Munich, Freising, Germany; Centre for Individualised Infection Medicine (CiiM), a joint venture between the Helmholtz Centre for Infection Research (HZI) and Hannover Medical School(MHH), Hannover, Germany; Department of Gastroenterology, Hepatology, Infectious Diseases and Endocrinology, Hannover Medical School (MHH), Hannover, Germany; TWINCORE, a joint venture between the Helmholtz-Centre for Infection Research (HZI) and the Hannover Medical School (MHH), Hannover, Germany; CAIMed – Center for AI in Medicine, Joint Venture of Leibniz University Hannover and Hannover Medical School, Hannover, Germany; Cluster of Excellence Resolving Infection Susceptibility (RESIST; EXC 2155), Hannover Medical School, Hannover, Germany; German Centre for Infection Research (DZIF), Partner Site Hannover-Braunschweig, Braunschweig, Germany; Munich Data Science Institute (MDSI), Technical University of Munich, 85748 Garching, Germany

**Keywords:** DNA methylation, RNA-seq, Software

## Abstract

**Summary:** Cell-type deconvolution is widely applied to gene expression and DNA methylation data, but access to methods for the latter remains limited. We introduce *deconvMe*, a new R package that simplifies access to DNA methylation-based deconvolution methods predominantly for blood data, and we additionally compare their estimates to those from gene expression and experimental ground truth data using a unique matched blood dataset.

## 1 Introduction

The human body consists of many cell types whose differentiation is characterized by changes in epigenomic and transcriptomic profiles. The exact composition of cell types in samples can provide valuable insights into, e.g., disease subtype or treatment response (Finotello and Trajanoski 2018). Experimental techniques to measure cell-type composition include flow cytometry and immunohistochemistry (IHC), which both are limited in their throughput and the number of possible cell types being profiled (Petitprez et al. 2018). The most cost-effective and most widely used assays for studying the transcriptome or epigenome, RNA sequencing (RNA-seq) (Wang, Gerstein, and Snyder 2009) and Illumina arrays (EPIC or 450k), respectively, produce bulk profiles where the exact cellular composition is unknown. A wide variety of deconvolution methods have been proposed for both gene expression and DNA methylation (DNAm) data. These use cell-type specific gene expression or DNAm markers to deconvolve bulk samples by solving linear equations (Merotto et al. 2023). Most rely on regression-based algorithms, such as non-negative least squares regression (NNLS) (Moss et al. 2018) or nu-support vector regression (nu-SVR) (Newman et al. 2015). Others use optimization and projection methods like quadratic programming under linear constraints (Houseman et al. 2012), as well as generative and latent variable models (Hicks and Irizarry 2019). As no single approach outperforms all others, running several approaches in parallel is often necessary. To this end, we have previously developed the immunedeconv package (Sturm et al. 2019) for first-generation transcriptomics deconvolution methods that rely on pre-built gene signatures, and omnideconv (Dietrich et al. 2024) for second-generation methods that dynamically learn cell-type specific gene signatures from single-cell expression data. While deconvolution methods for gene expression (Sturm et al. 2019; Dietrich et al. 2024) and DNAm-based methods (Teschendorff et al. 2017; De Ridder et al. 2024) have been extensively benchmarked, it is unclear which of the two data types offers more precise cell-type estimates. While selected methods of the two categories have been compared previously (Zheng et al. 2018), a dedicated performance comparison of methods and their signatures against a ground truth is currently lacking.

In this study, we present *deconvMe*, a novel R package that provides uniform access to five reference-based deconvolution methods (Suppl. Table 1) using DNAm array data: EpiDISH (Teschendorff et al. 2017), the Houseman method (Houseman et al. 2012), MethAtlas (Moss et al. 2018), methylCC (Hicks and Irizarry 2019), and MethylResolver (Arneson, Yang, and Wang 2020). The goal of this work is twofold: first, to introduce *deconvMe* as a well-documented, user-friendly, and consistent tool for DNAm-based cell-type deconvolution (Fig. 1), primarily targeting blood data; and second, to compare the performance of DNAm-based and RNA-seq-based deconvolution methods. To enable this comparison, we generated a unique dataset of 59 peripheral blood mononuclear cell (PBMC) samples with matched DNA methylation (EPIC array), RNA sequencing, and flow cytometry measurements (Fig. 1). The latter serve as ground truth to evaluate the accuracy of the respective deconvolution approaches.

**Figure 1:**
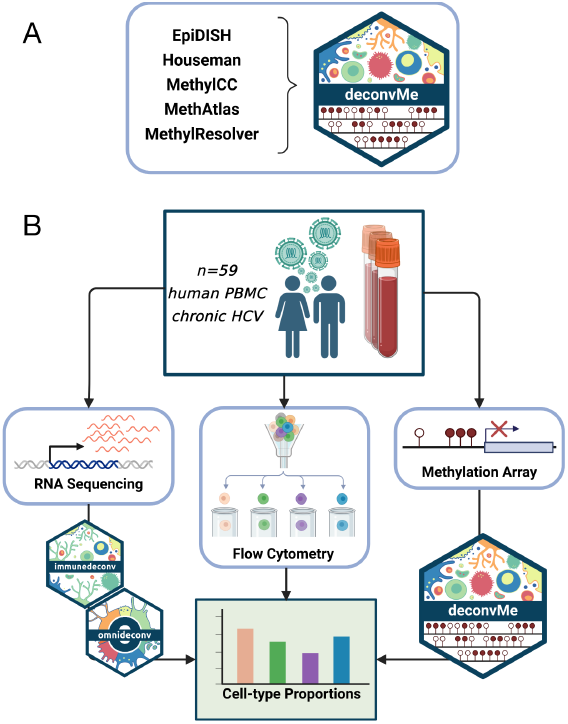
Methods included in *deconvMe* and comparison to RNA-seq based deconvolution. Five reference-based deconvolution methods are included in *deconvMe* (A), which we applied to our dataset of 59 chronic HCV patients with matched readouts for RNA-seq, EPIC DNAm arrays and flow-cytometry derived cell-type proportions. The RNA-seq samples were processed with deconvolution methods from immunedeconv and omnideconv and finally compared with ground truth cell-type proportions from flow cytometry (B).

## 2 Methods

### 2.1 Data Generation

We collected peripheral blood samples from 20 patients with chronic HCV infection at three time points: before treatment, after treatment with direct-acting antivirals, and at long-term follow-up. Peripheral blood mononuclear cells (PBMCs) were isolated using density centrifugation, cryopreserved, and later processed for quality control and downstream analysis. One sample had to be removed due to low quality. From each of the resulting 59 samples, we generated matched DNA methylation data using Illumina EPIC arrays, bulk RNA sequencing, and immune cell-type proportions via flow cytometry (Fig. 1). This matched multi-omic dataset enables direct comparison of deconvolution methods using a consistent biological ground truth. More details on the assays, the antibody panel, and data processing are available as supplementary material.

### 2.2 *deconvMe* implementation

We implemented *deconvMe* as an R package that uses a methylSet (Aryee et al. 2014) object containing the matrices for methylated and unmethylated probes as input. All methods we included are reference-based, due to their superiority compared to reference-free methods (Song and Kuan 2022). While all methods support blood data, support for other tissues is limited. Each method can easily be executed using deconvolute(), where the method parameter specifies which method should be executed. Additionally, we transform the output of each method to a unified style with samples in rows and estimated cell-type fractions in columns. Further, multiple methods can be executed at the same time, and their results aggregated. To directly access all parameters and results of the original method, we additionally provide functions called run_<methodname>. Each parameter is initially set to the method’s default values. Both MethylCC and the Houseman method can differentiate between the 450k and EPIC array types, a functionality that we also exposed to the user of *deconvMe*. Each method can be executed with a user-defined subset of features in their original signature and, finally, we provide access to several visualization functions. We further give direct access to each method’s signature matrix to inspect them further and compared how results change by exchanging them between methods, a feature natively supported by *deconvMe*. In our analysis we compared the genomic overlap of features in signature matrices of DNAm-based and gene expression-based methods. Details on these comparisons are available as supplementary material.

### 2.3 Deconvolution

Deconvolution methods using DNAm data as input were used with default parameters as implemented in *deconvMe*. For a comparison with gene expression-based methods, both first- and second-generation methods can be considered. For the first-generation methods, which come with pre-built cell-type specific signatures, we selected the three best-performing methods from the immunedeconv package (CIBERSORT (Newman et al. 2015), EPIC (Racle and Gfeller 2020), quanTIseq (Finotello et al. 2019)). Among second-generation methods, which generate their signatures dynamically from single-cell RNA-seq data, we selected the three best-performing methods from the omnideconv package (Bisque (Jew et al. 2020), DWLS (Tsoucas et al. 2019), and Scaden (Menden et al. 2020)) were trained with a single-cell RNA-seq (scRNA-seq) dataset from human blood (Hao et al. 2021), executed with default parameter settings and normalizations as proposed in omnideconv. Details on parameter settings, data generation, processing, and cell-type labelling (bulk RNA-seq, DNAm, and scRNA-seq) are available as supplementary material.

## 3 Application: DNAm vs gene-expression-based deconvolution

To illustrate the usability of *deconvMe*, we generated sample-matched readouts from chronic Hepatitis C virus-infected patients for DNAm using Illumina EPIC arrays, gene expression using RNA-seq (Suppl. Fig. 1) as well as proportions of selected immune cells using flow cytometry (Fig. 1). With this unique dataset, we could compare the results of reference-based cell-type deconvolution methods implemented in *deconvMe* to the matched set of ground-truth flow cytometric data and additionally evaluated their performance against gene-expression-based deconvolution methods included in immunedeconv and omnideconv. We selected five methods that can estimate immune cell proportions from DNAm data to be included in *deconvMe* and as a base of our comparison: MethAtlas, methylCC, MethylResolver, EpiDiSH and the Houseman method (Suppl. Table 1). All methods are reference-based, meaning they include a cell-type specific reference based on CpGs or - in the case of methylCC - differentially methylated regions (DMRs) based on prior knowledge. Similar types of methods with fixed references exist for gene expression measurements, such as from RNA-seq experiments. Beyond those, methods have been developed that are also designed for the deconvolution RNA-seq data, but can learn their references from scRNA-seq data. From both groups of methods, we, respectively, selected three of the currently best-performing ones (Sturm et al. 2019; Dietrich et al. 2024), and compared their results with the DNAm-based methods. As not all references are designed for the same sets of cell types, we only selected a set of representative immune cells that are estimated by almost every method (Suppl. Table 2).

We evaluated the performance using the ground truth from flow cytometric analysis with two metrics, Pearson’s R and root mean square error (RMSE). Of the DNAm-based methods, MethylCC, Houseman and MethylResolver perform similarly globally (Figure 2), followed by EpiDISH and MethAtlas. On a cell-type level, MethAtlas struggles with a correct estimation of CD8+ T cells, where it only reaches an RMSE of 0.2 due to overestimating this cell type in all samples; conversely, the same method underestimates CD4+ T cells (Suppl. Fig. 2). Similarly, while EpiDISH could perform well globally, as previously shown (De Ridder et al. 2024), it displays the weakest performance for NK cells.

**Figure 2:**
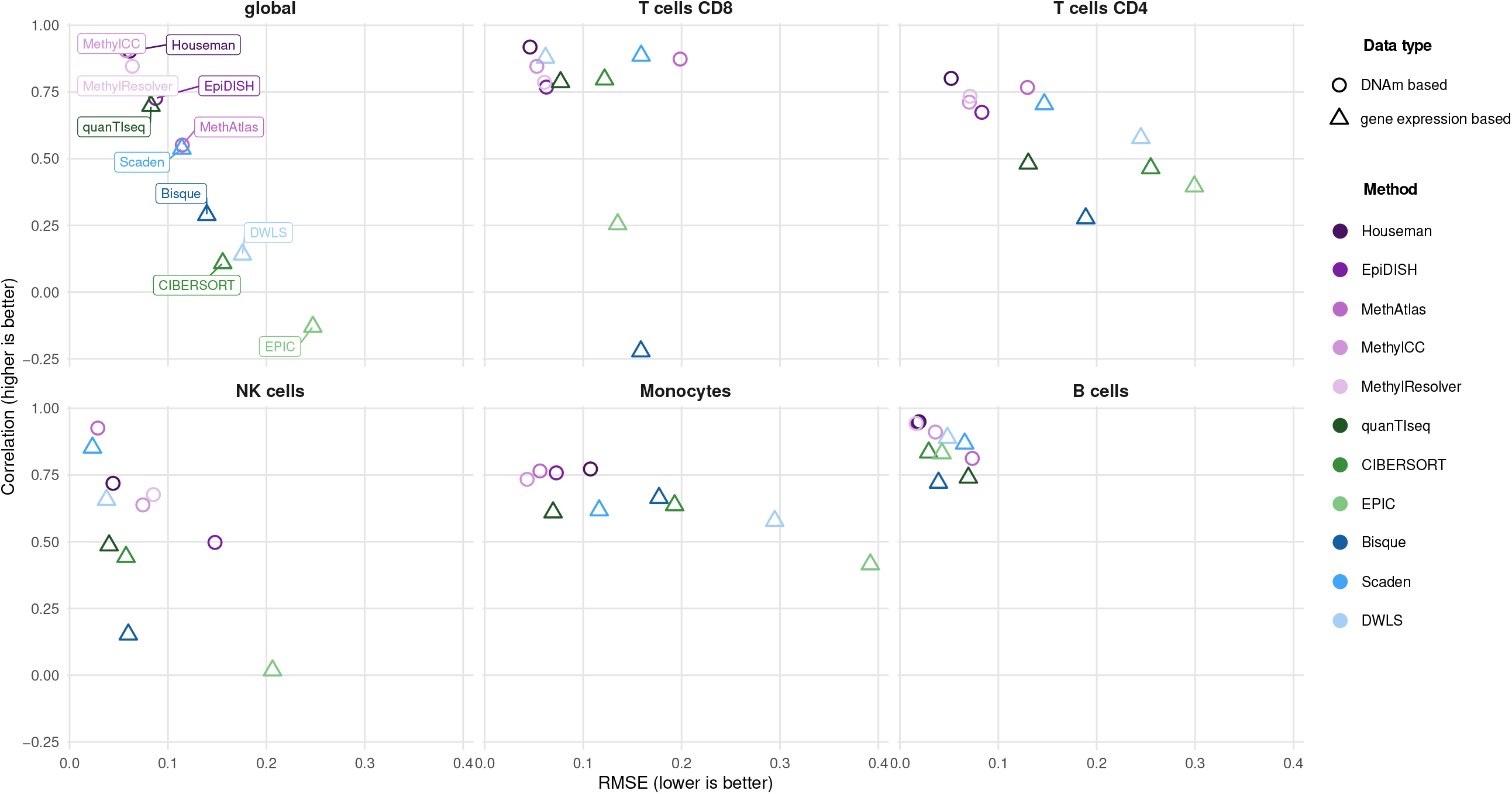
DNAm-based and gene expression-based deconvolution performances. Pearson correlation and RMSE for eleven deconvolution methods, based on flow-cytometry derived ground-truth fractions. Metrics are calculated globally (i.e., using the cell-type fractions from all cell types, displayed in the first box) and cell-type specifically. Methods can be grouped into two groups, based on their respective input data type (DNAm or gene expression readouts). Methods that use gene expression data can be further divided into first (green shades) and second (blue shades) generation methods, depending on the signature creation step. All DNAm-based methods (purple shades) are easily accessible via *deconvMe*.

Comparing the DNAm-based methods to those using gene-expression data (Fig. 2, Suppl. Fig. 2), we could spot systematic differences, as only quanTIseq and Scaden could match the performances of the *deconvMe*-included methods globally. While for B cells, all eleven methods had similar high performances, in other cell types the DNAm-based methods could consistently outperform gene-expression based methods on at least one performance metric. The single-cell informed methods hereby had no systematic performance increase compared to the first generation of gene-expression-based methods.

We further looked into the genomic overlaps of features included in the fixed signatures of the displayed deconvolution methods (Suppl. Fig. 3). We did not include the references from the single-cell informed deconvolution methods in this analysis, as they are highly reference data specific and therefore not directly comparable to the fixed references included in the other deconvolution methods. Strikingly, only a few CpGs are shared between the DNAm-based deconvolution methods, in the case of MethylCC, not a single position of its cell-type specific DMRs is covered by any other gene or CpG. As all methods performed similarly, this low overlap shows the variety of potential cell-type specific methylation sites in the human genome. This is confirmed further, as deconvolution performance still remains high when exchanging signature matrices for EpiDISH and MethAtlas (Suppl. Fig. 4). MethylResolver, however, shows a higher dependency on its original signature, as its performance drops once signatures from EpiDISH, Houseman, or MethAtlas are used. Conversely, genes are more commonly shared between signature matrices, although most are unique to a single method. Interestingly, almost no overlaps are present between the two data types, hinting at different layers of cell-type specificity in these data types.

## 4 Conclusion

Despite the increasing popularity of single-cell techniques, bulk profiling is still common for large-scale projects due to the low costs and well-established workflows. Moreover, single-cell profiling is rarely used to study DNA methylation. Cell-type deconvolution methods overcome a limitation of bulk profiling by robustly estimating cell-type fractions. Here, we describe a novel R package *deconvMe*, that offers simplified access to five cell-type deconvolution methods tailored for DNAm data. We compared the performance of these methods to RNA-seq based cell-type deconvolution methods and show that they perform better in terms of RMSE and Pearson correlation compared to gene-expression-based methods. Importantly, *deconvMe* allows users to run multiple DNAm-based methods in parallel - many of which have shown similar performance across prior benchmarks - and compare their results side by side, while allowing methods to additionally accept external signature matrices. This is particularly valuable in settings where no method clearly outperforms the others. To support this benchmarking, we provide access to a matched DNA methylation (EPIC array), RNA-seq, and flow cytometry dataset. To our knowledge, no comparable dataset exists, making it a valuable resource for future method development and benchmarking in the deconvolution field. While flow cytometry serves as a practical ground truth, it has limitations, particularly for rare or hard-to-measure cell types (Liu et al. 2024). Pseudobulk profiles from single-cell data may serve as an alternative, but such data are still scarce for DNAm, especially in complex tissues. Together, *deconvMe* and our dataset aim to make DNAm-based deconvolution methods more accessible and comparable, supporting future benchmarking efforts and broader adoption in the research community.

## Author Contributions

AD and ML devised the concept for *deconvMe*. AD and KP created the initial version of *deconvMe*. LLW continued this work until its completion and helped run the deconvolution methods. AD did the remaining analysis steps and package finalizing. ML and AD provided supervision of KP and LLW. CO initiated and coordinated the database and laboratory work. MW and SK acquired the laboratory data, supervised by AK and MC. AD drafted the manuscript. All authors revised the manuscript and gave their approval.

## Supplementary data

Supplementary data are available at *Bioinformatics Advances* online.

## Conflict of interest

MC reports personal fees from Abbvie, personal fees from Falk Foundation, personal fees from Gilead, personal fees from GlaxoSmithKline, personal fees from Jansen-Cilag, personal fees from Merck/MSD, personal fees from Novartis, personal fees from Roche, personal fees from Spring Bank Pharmaceuticals, and personal fees from Swedish Orphan Biovitrum, outside the submitted work. ML consults for mbiomics GmbH. Other authors have nothing to disclose.

## Funding

This work was supported by the German Federal Ministry of Education and Research (BMBF) within the framework of the CompLS research and funding concept [031L0294 (NetfLID)].

## Software and Data availability

*deconvMe* is available at https://github.com/omnideconv/deconvMe. The scripts to conduct the analysis steps and figures can be found at https://github.com/omnideconv/deconvMe_analysis. Matched EPIC array data, RNA-seq data, and flow cytometry values are available at https://doi.org/10.6084/m9.figshare.28563854.v2.

## Notes

https://github.com/omnideconv/deconvMe

https://doi.org/10.6084/m9.figshare.28563854.v2

### Summary of Updates

We renamed the package, added comparisons to scRNA-seq based deconvolution methods and improved our Figure 1. Also, we added new supply. plots that show how deconvolution performance changes when signatures from different methods are used.

