## Supplementary for "Unifying DNA methylation-based in silico cell-type deconvolution with *deconvMe*"

### Supplementary Figures

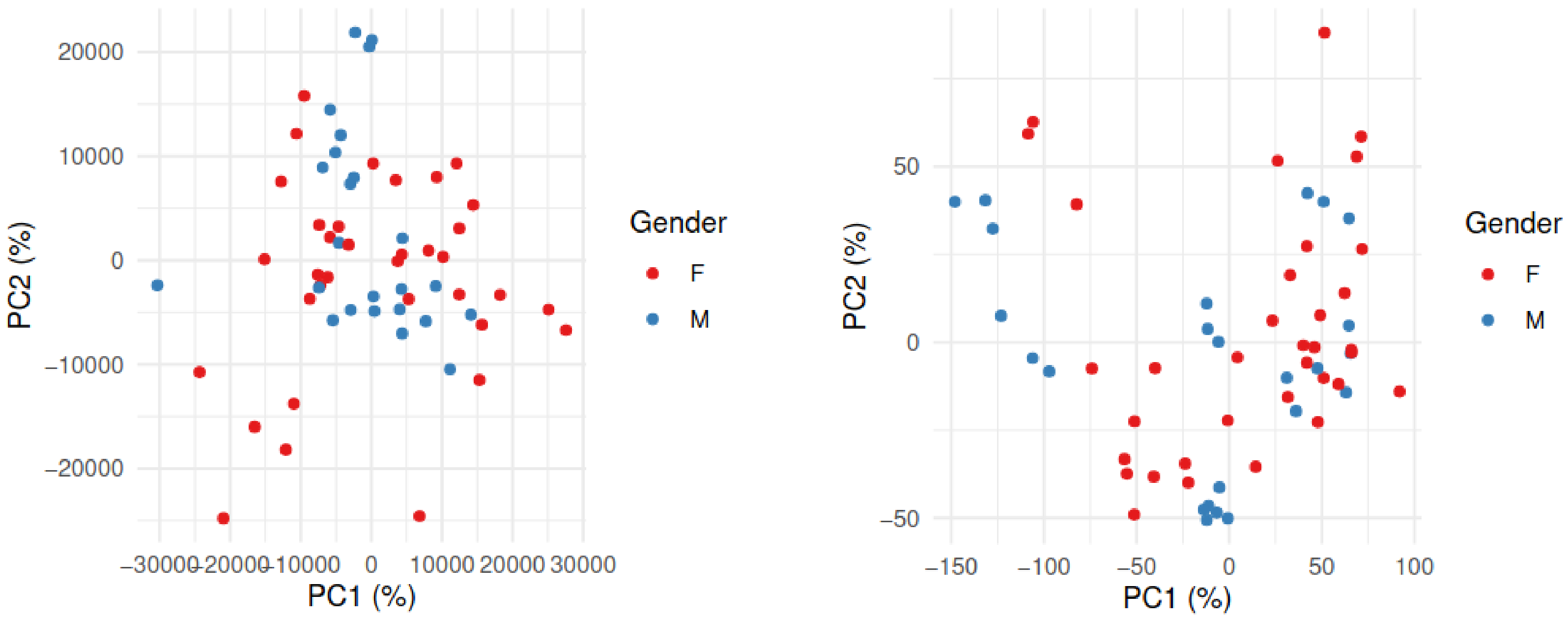

**Supplementary Figure 1:** Principal component analysis (PCA) shows the first two PCs for DNAm data (right) and RNA-seq data (left) of the same samples (n=59).

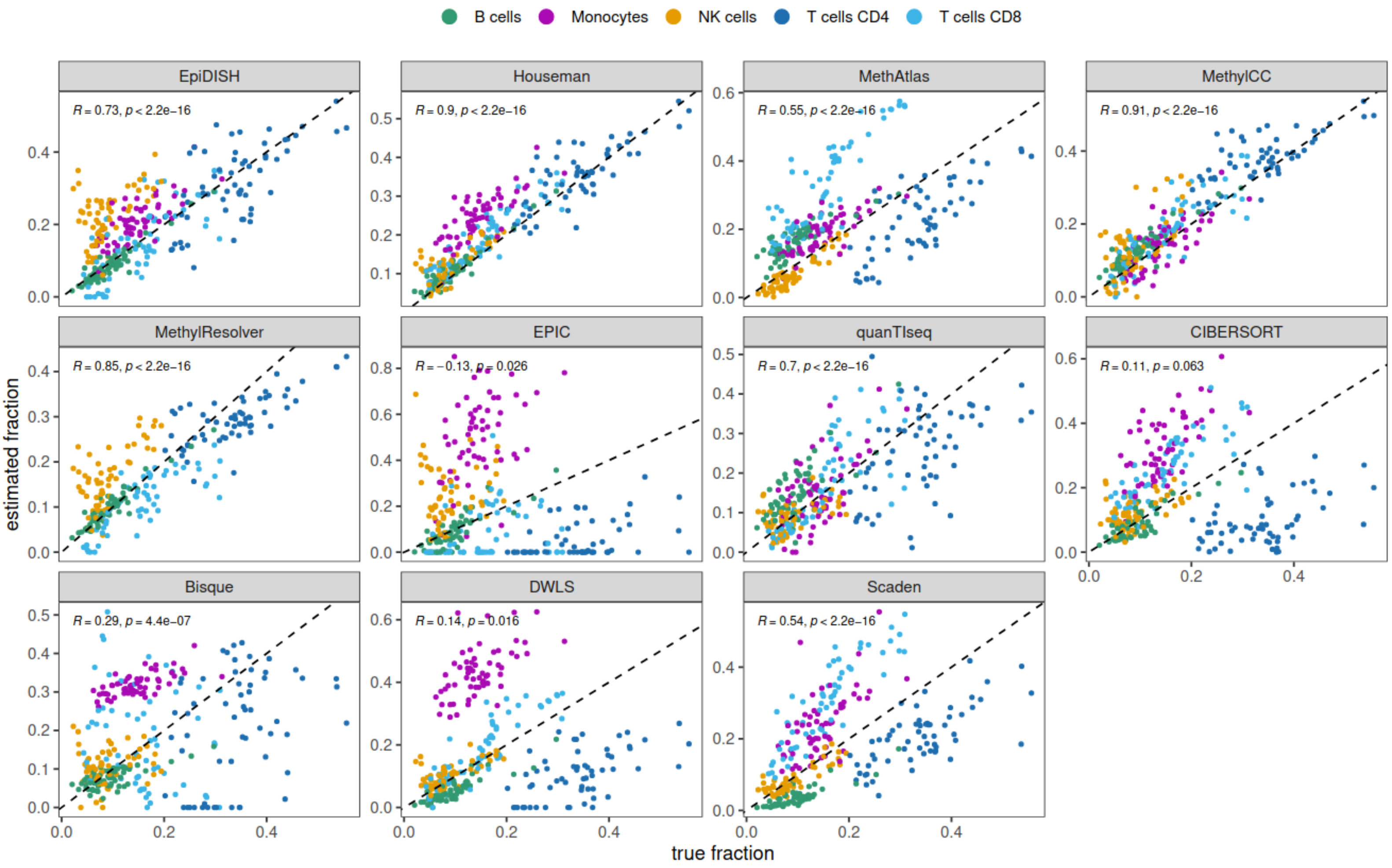

**Supplementary Figure 2:** Pearson correlation of cell-type fractions predicted by deconvolution methods versus ground truth in 59 samples from bulk DNAm (EpiDISH, Houseman, MethAtlas, MethylCC, MethylResolver) and RNA-seq (EPIC, quanTIseq, CIBERSORT, Bisque, DWLS, Scaden). The ground truth is derived from flow cytometry.

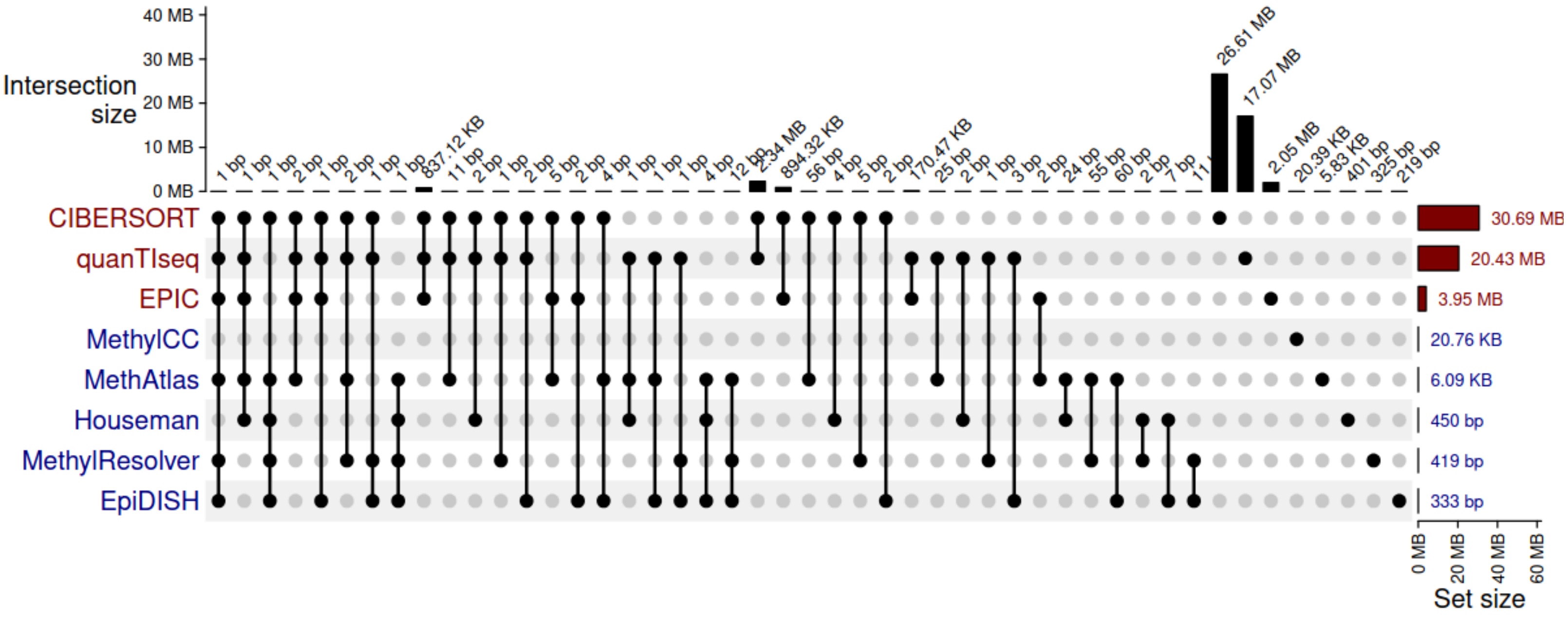

**Supplementary Figure 3:** Upset plot of signature matrix features of eight deconvolution methods (gene-expression-based in red, DNAm-based in blue) showing the overlap of genomic positions between those features. The horizontal barplot at the right shows the size of all features in the sum of base pairs (one CpG is counted as a single base pair in this setting). Vertical bar plots at the top show the size of intersections in the number of base pairs. Regions of genes include exons, introns, and promoters.

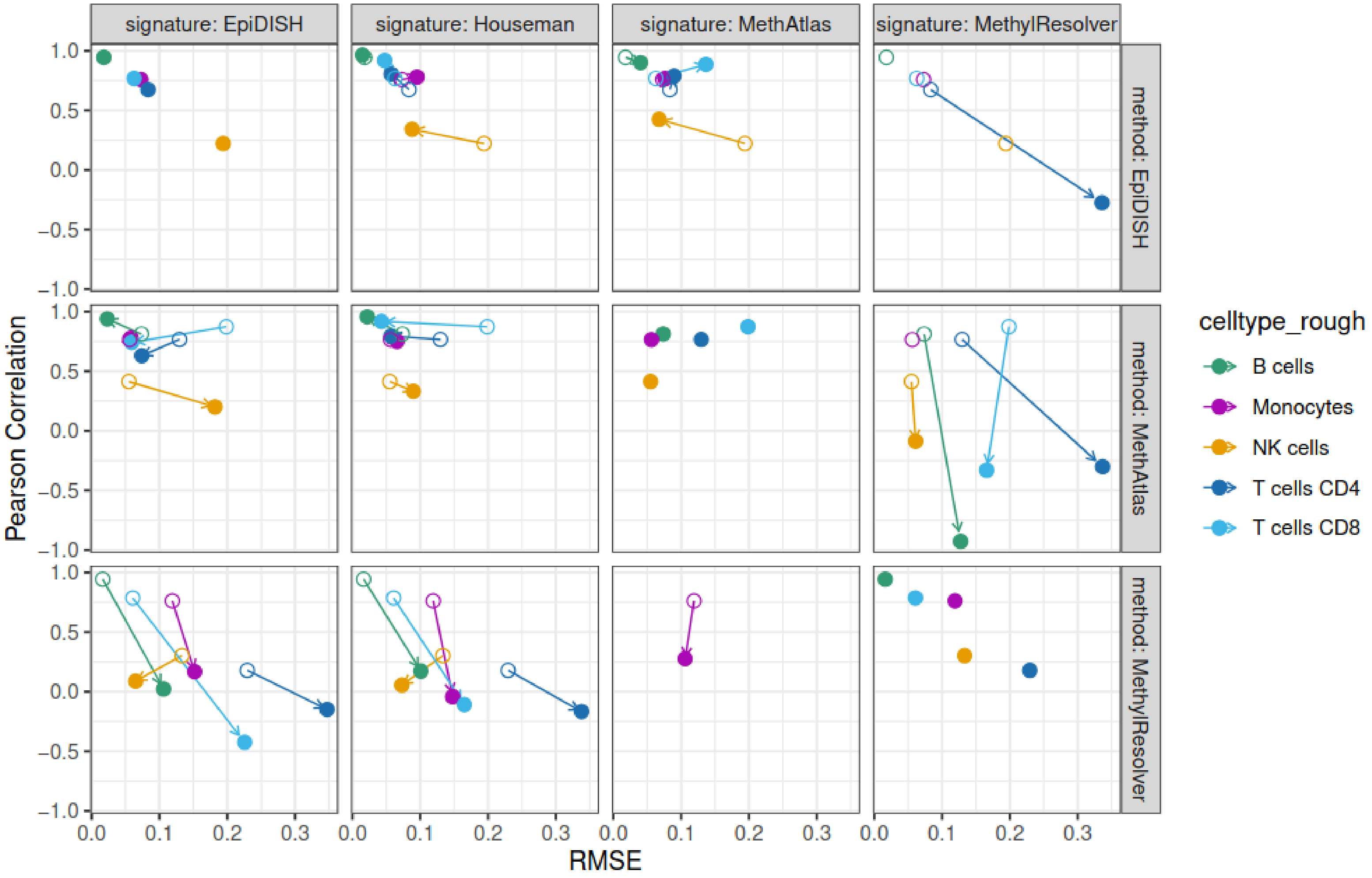

**Supplementary Figure 4:** Pearson correlation and RMSE (as for Figure 2) for cell type estimates when calculated by different combinations of DNAm-based deconvolution algorithm (rows) and signature matrices (columns). Filled-out points show the actual results of the indicated method-signature combination, while open points show the baseline result with the original method and its corresponding signature. The arrows indicate the change in correlation and RMSE due to a change in signature, while keeping the method fixed.

The HCV PBMC dataset was used, and cell types were relabelled as described in Suppl. Table 2. Missing data points indicate that the cell type was not estimated in any sample by a method+signature combination, and the correlation coefficient can not be calculated.

### Supplementary Tables

| **Method** | **Algorithm** | **Intended Tissue Type(s)** | **Signature Source Data** | **Reference** |
|---|---|---|---|---|
| EpiDISH | Robust Partial Correlations (RPC), Constrained Projection (CP), Support Vector Regression (SVR) | Blood, Epithelial cells, Breast | purified blood, epithelial, and non-epithelial data, extended with DNAse Hypersensitivity Sites from Roadmap and ENCODE | (Teschendorff et al. 2017) |
| Houseman | Constrained projection/Quadratic Programming (CP/QP) | Blood | Purified blood cell methylation profiles (IDOL-optimized) | (Houseman et al. 2012; Koestler et al. 2016) |
| MethylCC | Constrained Linear model based on DMRs | Blood, can be extended to other tissues | purified blood | (Hicks and Irizarry 2019) |
| MethylResolver | Least Trimmed Squares (LTS) regression | Blood, cancer tissue | Purified blood methylation profiles (IDOL-optimized), extended with additional leukocytes | (Arneson, Yang, and Wang 2020) |
| MethAtlas | non-negative least squares (NNLS) | Blood, immune, tissue-wide | Comprehensive atlas of purified tissue and immune cell methylomes | (Moss et al. 2018) |

**Supplementary Table 1:** Reference-based methods included in *deconvMe*. For each method, the core algorithm that solves the deconvolution problem is listed; further, the intended tissue(s) for which the method was implemented or tested on, and the dataset types that were used to create the internal signature matrices.

| celltype_coarse | celltype_clean | celltype_original | method |
|---|---|---|---|
| B cells | B cells | B | EpiDISH |
| T cells CD4 | T cells CD4 | CD4T | EpiDISH |
| T cells CD8 | T cells CD8 | CD8T | EpiDISH |
| other | Eosinophils | Eosino | EpiDISH |
| Monocytes | Monocytes | Mono | EpiDISH |
| other | Neutrophils | Neutro | EpiDISH |
| NK cells | NK cells | NK | EpiDISH |
| B cells | B cells | Bcell | Houseman |
| T cells CD4 | T cells CD4 | CD4T | Houseman |
| T cells CD8 | T cells CD8 | CD8T | Houseman |
| Monocytes | Monocytes | Mono | Houseman |
| other | Neutrophils | Neu | Houseman |
| NK cells | NK cells | NK | Houseman |
| B cells | B cells | B-cells_EPIC | MethAtlas |
| T cells CD4 | T cells CD4 | CD4T-cells_EPIC | MethAtlas |
| T cells CD8 | T cells CD8 | CD8T-cells_EPIC | MethAtlas |
| Monocytes | Monocytes | Monocytes_EPIC | MethAtlas |
| other | Neutrophils | Neutrophils_EPIC | MethAtlas |
| NK cells | NK cells | NK-cells_EPIC | MethAtlas |
| B cells | B cells | Bcell | MethylCC |
| T cells CD4 | T cells CD4 | CD4T | MethylCC |
| T cells CD8 | T cells CD8 | CD8T | MethylCC |
| other | Granulocytes | Gran | MethylCC |
| Monocytes | Monocytes | Mono | MethylCC |
| NK cells | NK cells | NK | MethylCC |
| B cells | B cells | Bcell | MethylResolver |
| T cells CD8 | T cells CD8 | CD8 | MethylResolver |
| other | Dendritic cells | Dendritic | MethylResolver |
| other | Eosinophils | Eos | MethylResolver |
| other | Macrophages | Macro | MethylResolver |
| other | other | Mon | MethylResolver |
| other | Neutrophils | Neu | MethylResolver |
| NK cells | NK cells | NK | MethylResolver |
| T cells CD4 | T cells CD4 | Tmem | MethylResolver |
| T cells CD4 | T cells CD4 | Tnaive | MethylResolver |

| | | | |
|---|---|---|---|
| other | Tregs | Treg | MethylResolver |
| B cells | B cells | B cell | quanTIseq |
| other | Macrophages | Macrophage M1 | quanTIseq |
| other | Macrophages | Macrophage M2 | quanTIseq |
| Monocytes | Monocytes | Monocyte | quanTIseq |
| other | Neutrophils | Neutrophil | quanTIseq |
| NK cells | NK cells | NK cell | quanTIseq |
| T cells CD4 | T cells CD4 | T cell CD4+ (non-regulatory) | quanTIseq |
| T cells CD8 | T cells CD8 | T cell CD8+ | quanTIseq |
| other | Tregs | T cell regulatory (Tregs) | quanTIseq |
| other | Dendritic cells | Myeloid dendritic cell | quanTIseq |
| other | other | uncharacterized cell | quanTIseq |
| B cells | B cells | B cell | EPIC |
| T cells CD4 | T cells CD4 | T cell CD4+ | EPIC |
| T cells CD8 | T cells CD8 | T cell CD8+ | EPIC |
| Monocytes | Monocytes | Monocyte | EPIC |
| NK cells | NK cells | NK cell | EPIC |
| other | Neutrophils | Neutrophil | EPIC |
| other | other | uncharacterized cell | EPIC |
| B cells | B cells | B cell naive | CIBERSORT |
| B cells | B cells | B cell memory | CIBERSORT |
| B cells | B cells | B cell plasma | CIBERSORT |
| T cells CD8 | T cells CD8 | T cell CD8+ | CIBERSORT |
| T cells CD4 | T cells CD4 | T cell CD4+ naive | CIBERSORT |
| T cells CD4 | T cells CD4 | T cell CD4+ memory resting | CIBERSORT |
| T cells CD4 | T cells CD4 | T cell CD4+ memory activated | CIBERSORT |
| T cells CD4 | T cells CD4 | T cell follicular helper | CIBERSORT |
| other | Tregs | T cell regulatory (Tregs) | CIBERSORT |
| other | gd T cells | T cell gamma delta | CIBERSORT |
| NK cells | NK cells | NK cell resting | CIBERSORT |
| NK cells | NK cells | NK cell activated | CIBERSORT |
| Monocytes | Monocytes | Monocyte | CIBERSORT |
| other | Macrophages | Macrophage M0 | CIBERSORT |
| other | Macrophages | Macrophage M1 | CIBERSORT |
| other | Macrophages | Macrophage M2 | CIBERSORT |
| other | Dendritic cells | Myeloid dendritic cell resting | CIBERSORT |
| other | Dendritic cells | Myeloid dendritic cell activated | CIBERSORT |

| | | | |
|---|---|---|---|
| other | other | Mast cell activated | CIBERSORT |
| other | other | Mast cell resting | CIBERSORT |
| other | Eosinophils | Eosinophil | CIBERSORT |
| other | Neutrophils | Neutrophil | CIBERSORT |
| B cells | B cells | B cells | DWLS |
| other | Innate lymphoid cells | ILC | DWLS |
| other | Dendritic cells | mDC | DWLS |
| Monocytes | Monocytes | Monocytes | DWLS |
| NK cells | NK cells | NK cells | DWLS |
| other | Plasmacytoid dendritic cells | pDC | DWLS |
| other | Plasma cells | Plasma cells | DWLS |
| other | Platelets | Platelet | DWLS |
| T cells CD4 | T cells CD4 | T cells CD4 conv | DWLS |
| T cells CD8 | T cells CD8 | T cells CD8 | DWLS |
| other | Tregs | Tregs | DWLS |
| B cells | B cells | B cells | Scaden |
| other | Innate lymphoid cells | ILC | Scaden |
| other | Dendritic cells | mDC | Scaden |
| Monocytes | Monocytes | Monocytes | Scaden |
| NK cells | NK cells | NK cells | Scaden |
| other | Plasmacytoid dendritic cells | pDC | Scaden |
| other | Plasma cells | Plasma cells | Scaden |
| other | Platelets | Platelet | Scaden |
| T cells CD4 | T cells CD4 | T cells CD4 conv | Scaden |
| T cells CD8 | T cells CD8 | T cells CD8 | Scaden |
| other | Tregs | Tregs | Scaden |
| B cells | B cells | B cells | Bisque |
| other | Innate lymphoid cells | ILC | Bisque |
| other | Dendritic cells | mDC | Bisque |
| Monocytes | Monocytes | Monocytes | Bisque |
| NK cells | NK cells | NK cells | Bisque |
| other | Plasmacytoid dendritic cells | pDC | Bisque |
| other | Plasma cells | Plasma cells | Bisque |

| | | | |
|---|---|---|---|
| other | Platelets | Platelet | Bisque |
| T cells CD4 | T cells CD4 | T cells CD4 conv | Bisque |
| T cells CD8 | T cells CD8 | T cells CD8 | Bisque |
| other | Tregs | Tregs | Bisque |
| B cells | B cells | CD19+ (B-Zellen) | FACS |
| Monocytes | Monocytes | CD14+ (Monozyten) | FACS |
| other | MAIT cells | MAIT Zellen | FACS |
| other | gd T cells | gd T-Zellen | FACS |
| other | other | CD8+, CD4+ | FACS |
| other | other | CD8-, CD4- | FACS |
| T cells CD4 | T cells CD4 | T- Helferzellen | FACS |
| T cells CD8 | T cells CD8 | zytotoxische T-Zellen | FACS |
| other | Dendritic cells | DCs | FACS |
| other | other | non DCs | FACS |
| NK cells | NK cells bright | NK bright | FACS |
| NK cells | NK cells dim | NK dim | FACS |
| other | other | non NK | FACS |
| other | other | non LD- | FACS |

**Supplementary Table 2:** Cell-type mapping that was performed to ensure comparability between methods in case of different spelling or annotation levels. The first three columns represent the aggregated cell-type labels on the highest cell-type level (celltype_coarse), the original cell-type labels renamed to a controlled vocabulary (celltype_clean) and the original cell-type labels of each method (celltype_original), respectively.

#### Data Generation and Processing

We generated all data based on a cohort of 20 patients chronically infected with the Hepatitis C virus. Patients were recruited at the Hannover Medical School outpatient clinic between 1 January 2014 and 31 December 2020. Blood was drawn before treatment, after treatment with direct-acting antivirals (DAA), and at long-term follow-up (48 or 96 weeks after treatment), allowing us to study different phases of infection. The cohort was extensively characterized, and samples were prospectively collected and stored in a biobank. In short, blood was drawn, and peripheral blood mononuclear cells (PBMCs) were isolated according to a standard Ficoll Hypaque density centrifugation protocol (BioColl separating solution; Biochrom AG, Berlin, Germany). After isolation, cells were transferred to a freezing medium and stored in liquid nitrogen. For analysis purposes, the cells were thawed, counted, and then further processed.

DNA methylation has been measured using Illumina EPIC arrays. DNA isolation was conducted using the Monarch Genomic DNA Purification Kit T3010L (New England Biolabs, Ipswich, MA, USA) according to the manufacturer's instructions. We normalized the DNA concentration of all samples after elution to 50 ng/µl, randomized the samples on a 96-well plate. We stored the plate at -80 °C. DNAm measurement was performed at the Human Genomics Facility of Erasmus MC, Rotterdam, the Netherlands. 500 ng of DNA was bisulfite converted using the EZ-96 DNA Methylation kit (Zymo Research Corp., Irvine, CA, USA) with the KingFisher Flex robot (Thermo Fisher Scientific, Breda, the Netherlands). The methylation status of 59 samples was assessed in 8 µl bisulfite-treated DNA using the Infinium MethylationEPIC BeadChip, following Illumina's protocol.

Gene expression has been measured using bulk RNA sequencing. For Bulk-RNA sequencing, one million PBMCs were thawed, and cells were directly resuspended in 700 µl RLT buffer (Qiagen). RNA extraction and sequencing library synthesis followed the manufacturer's protocol (RNeasy plus microRNA extraction kit; Qiagen and Single-cell low Input Kit, NEB). Subsequently, RNA samples were sequenced at HZI Braunschweig with an Illumina NovaSeqS2 (100 cycles), aiming for 50 million reads per sample.

Additionally, samples were subjected to flow cytometry (FACS). Antibodies: FITC, PE, PE-Cy7, APC-R700, APC-H7, BV510, BV605, BV711, BV786 (BD, Franklin Lakes, NJ, USA and Biolegend, San Diego, CA, USA) were used to determine cell types. Our flow-cytometric-gating filtered for lymphocytes and single cells and excluded dead cells (LD+). Monocytes (CD14+), B cells (CD19+), CD4+ T cells (CD19-, CD3+, CD4+, CD8-), CD8+ T cells (CD19-, CD3+, CD4-, CD8+) and NK cells (CD19-, CD3-, CD56+) were used in our analysis. 1 sample was excluded since the flow data did not pass our internal quality control. Overall, we included 59 samples in our analysis.

Raw fastq files from the RNA sequencing have been processed using the nf-core rnaseq pipeline version 3.10.1 (Patel et al. 2024; Ewels et al. 2020). The main steps include a manual inspection of quality using FastQC, followed by pseudo-alignment and gene quantification using Salmon (Patro et al. 2017). Next to raw gene expression counts, this method generates transcript per million (TPM) counts, which are normalized by gene length and library size and were used as input for deconvolution methods. DNAm data by Illumina EPIC arrays generated raw idat files that have been processed with RnBeads 2.0 (Müller et al. 2019) using the default settings during preprocessing, which removes methylation sites with too many missing values, cross-reactive probes, or those that overlap with known single-nucleotide polymorphisms (SPNs). Additionally, we removed all CpGs with NA values in any sample. The final dataset consisted of 724,058 CpGs. We removed CpGs located on sex chromosomes as they would confound the analysis. We also removed one sample due to low quality in the RNA-seq and DNAm assays and had a final dataset of 59 samples.

### Parameter settings for deconvolution methods

#### DNAm-based methods

The five DNAm-based deconvolution methods included in *deconvMe* (EpiDISH, Houseman, MethAtlas, MethylCC, MethylResolver) were executed with the following parameters (for all five methods, un-normalized beta value matrices were used as input):
EpiDISH was executed with the RPC algorithm, the supplied reference for blood, and 100 maximum iterations. The Houseman method used the IlluminaHumanMethylationEPIC reference platform, the composite cell type was set to blood, probe selection was performed with the IDOL (Koestler et al. 2016) algorithm and internal preprocessing was done with preprocessQuantile. The signature matrix of MethAtlas was subset to only contain references for EPIC array data. MethylCC was set to expect EPIC array data, the epsilon parameter set to 0.01, max_iter to 100, and used the 'random' method to initialize parameter estimates. Finally, MethylResolver was run with an alpha value of 1 and in absolute mode.
All five methods were accessed via *deconvMe* version 1.1.0.

#### Gene-expression based methods

The first-generation methods used in this study (CIBERSORT, EPIC, and quanTIseq), were executed with the following parameters:
CIBERSORT was executed with TPM normalized expression values, the 'array' parameter to FALSE, and not in absolute mode. EPIC also used TPM normalized data, 'tumor' set to FALSE and scaling of mRNA switched on. Finally, quanTIseq also used TPM data, 'array' and 'tumor' to FALSE and scaling of mRNA switched on.
All three methods were accessed via *immunedeconv (Sturm et al. 2019)* version 2.1.0.

The second-generation methods (Bisque, DWLS, and Scaden) were executed as follows:
Bisque used TPM normalized RNA-seq data and CPM normalized counts for the scRNA-seq data; batches were defined by the sample IDs in the Hao scRNA-seq dataset. No marker genes were supplied, no overlapping samples between bulk and reference were expected.
DWLS and Scaden both used TPM normalized RNA-seq data and un-normalized counts for the scRNA-seq data. The signature of DWLS was built with a differential expression fold-change cutoff of 0.5 and p-value of 0.05, using the 'mast_optimized' algorithm. The deconvolution step further was executed with the 'DampenedWLS' algorithm.
Scadens signature was built with a batch size of 128, learning rate of 1e-4, 5000 steps, and 100 cells and 1000 samples for the simulation step.The follow-up deconvolution part has no exposed parameters.
All three methods were accessed via *omnideconv (Dietrich et al. 2024)* version 0.1.1.

### Single-cell RNA-seq data processing

We used a scRNA-seq dataset from human blood from (Hao et al. 2021) as reference dataset for three second-generation deconvolution methods. This CITE-seq dataset has been prepared previously by us for its dedicated usage in benchmarking such deconvolution methods in a recent work (Dietrich et al. 2024). Briefly, the dataset was filtered to remove low quality cells based on the number of expressed genes, number of counts per cell and mitochondrial content. Then, it was integrated with other blood scRNA-seq datasets to establish a combined, robust cell-type annotation, based on manual inspection of marker genes both on the cell surface and their RNA expression. The dataset was accessed from *deconvData* at https://doi.org/10.6084/m9.figshare.25348051.v2.

#### Cross-referencing methods and exchanging signature matrices

Through *deconvMe,* users are enabled to run supported methods also with external signature matrices instead of their original ones. This feature is possible for EpiDISH, MethylResolver, and MethAtlas. MethylCC has no signature matrix in the classical sense, as it works with differentially methylated regions (DMRs), and while the signature of the Houseman method is accessible, it cannot be replaced easily without breaking the method's structure.
We used this functionality to compare how deconvolution performance would change for EpiDISH, MethylResolver, and MethAtlas when using their original signatures compared to those of the respective other methods (Suppl. Fig. 4). Again, we used our PBMC dataset and calculated Pearson's R and RMSE using the flow-cytometry derived ground truth. MethylResolver drops in performance once external signatures are used, and in return, its signature causes EpiDISH and MethAtlas to underestimate and miss all cell types. The signatures of EpiDISH, Houseman, and MethAtlas only slightly change the deconvolution accuracy of EpiDISH and MethAtlas.

#### Comparison of signature matrices

We further compared the genomic overlap of features in the signature matrices of eight deconvolution methods. Signature matrices were extracted from the Github repositories or the respective package data. MethylCC uses differentially methylated regions (DMRs) instead of methylation sites as its signature features, which are calculated from the FlowSorted. Blood.450k whole blood reference methylomes for six cell types (Reinius et al. 2012; Jaffe et al. 2012) using the find_dmrs() function. In the following, the CpGs that are part of these DMRs are used for methylCC. To compare the genomic locations of all features on the same reference genome coordinates, we used the liftOver() function of the rtracklayer package (Lawrence, Gentleman, and Carey 2009; Hinrichs et al. 2006) in R to lift over all coordinates to the hg38 genome. This affected all DNAm-based methods, as the Illumina manifest files in which coordinates of 450k and EPIC probes are only available for hg19. The coordinates of genes in the reference matrices of quanTIseq, EPIC, and CIBERSORT were taken from the hg38 reference genome; this includes all exon, intron, and promoter regions of a gene.

#### Cell-type annotation

Following deconvolution, cell types were relabelled to allow comparisons between methods that either differed in the spelling of cell types or provided different annotation resolutions. In the latter case, estimates were summed up to the next ‘higher’ matching cell type (Supplementary Table 1).

Houseman, Eugene Andres, William P. Accomando, Devin C. Koestler, Brock C. Christensen, Carmen J. Marsit, Heather H. Nelson, John K. Wiencke, and Karl T. Kelsey. 2012. “DNA Methylation Arrays as Surrogate Measures of Cell Mixture Distribution.” *BMC Bioinformatics* 13 (May):86.

Jaffe, Andrew E., Andrew P. Feinberg, Rafael A. Irizarry, and Jeffrey T. Leek. 2012. “Significance Analysis and Statistical Dissection of Variably Methylated Regions.” *Biostatistics* 13 (1): 166–78.

Koestler, Devin C., Meaghan J. Jones, Joseph Usset, Brock C. Christensen, Rondi A. Butler, Michael S. Kobor, John K. Wiencke, and Karl T. Kelsey. 2016. “Improving Cell Mixture Deconvolution by Identifying Optimal DNA Methylation Libraries (IDOL).” *BMC Bioinformatics* 17 (March):120.

Lawrence, Michael, Robert Gentleman, and Vincent Carey. 2009. “Rtracklayer: An R Package for Interfacing with Genome Browsers.” *Bioinformatics (Oxford, England)* 25 (14): 1841–42.

Moss, Joshua, Judith Magenheim, Daniel Neiman, Hai Zemmour, Netanel Loyfer, Amit Korach, Yaacov Samet, et al. 2018. “Comprehensive Human Cell-Type Methylation Atlas Reveals Origins of Circulating Cell-Free DNA in Health and Disease.” *Nature Communications* 9 (1): 5068.

Müller, Fabian, Michael Scherer, Yassen Assenov, Pavlo Lutsik, Jörn Walter, Thomas Lengauer, and Christoph Bock. 2019. “RnBeads 2.0: Comprehensive Analysis of DNA Methylation Data.” *Genome Biology* 20 (1): 55.

Patel, Harshil, Phil Ewels, Jonathan Manning, Maxime U. Garcia, Alexander Peltzer, Rickard Hammarén, Olga Botvinnik, et al. 2024. *Nf-Core/rnaseq: Nf-Core/rnaseq v3.18.0 - Lithium Lynx*. Zenodo. https://doi.org/10.5281/ZENODO.14537300.

Patro, Rob, Geet Duggal, Michael I. Love, Rafael A. Irizarry, and Carl Kingsford. 2017. “Salmon Provides Fast and Bias-Aware Quantification of Transcript Expression.” *Nature Methods* 14 (4): 417–19.

Reinius, Lovisa E., Nathalie Acevedo, Maaike Joerink, Göran Pershagen, Sven-Erik Dahlén, Dario Greco, Cilla Söderhäll, Annika Scheynius, and Juha Kere. 2012. “Differential DNA Methylation in Purified Human Blood Cells: Implications for Cell Lineage and Studies on Disease Susceptibility.” *PloS One* 7 (7): e41361.

Sturm, Gregor, Francesca Finotello, Florent Petitprez, Jitao David Zhang, Jan Baumbach, Wolf H. Fridman, Markus List, and Tatsiana Aneichyk. 2019. “Comprehensive Evaluation of Transcriptome-Based Cell-Type Quantification Methods for Immuno-Oncology.” *Bioinformatics* 35 (14): i436–45.

Teschendorff, Andrew E., Charles E. Breeze, Shijie C. Zheng, and Stephan Beck. 2017. “A Comparison of Reference-Based Algorithms for Correcting Cell-Type Heterogeneity in Epigenome-Wide Association Studies.” *BMC Bioinformatics* 18 (1): 105.
